# Membrane remodeling by the Bacterial Dynamin-like Protein (BDLP) from *Nostoc punctiforme*

**DOI:** 10.1101/2025.05.27.656310

**Authors:** Keerti Singh, Thomas J. Pucadyil

**Affiliations:** Indian Institute of Science Education and Research, Dr. Homi Bhabha Road, Pashan, Pune 411008, Maharashtra, India

## Abstract

Membranes are fundamental to biological systems as they serve as barriers to compartmentalize the cell and its cytoplasm. They are constantly remodeled in shape and composition during the formation and maintenance of organelles. The large GTPase dynamins are well characterized for membrane shape remodeling in eukaryotes but their functions in prokaryotes are less characterized. Here, we determine the lipid-binding, enzymatic and membrane remodeling activities of the Bacterial Dynamin-Like Protein (BDLP) from the cyanobacterium *Nostoc punctiforme*. We find that BDLP binds phosphatidyl glycerol (PG)-containing membranes, but unlike other soluble dynamins, shows little stimulation in GTPase activity. This enzymatic feature is like that seen among the fusion dynamins to which BDLP shows structural similarities. FRET-based bulk vesicle fusion assays however reveal that BDLP is incapable of membrane fusion. To further understand BDLP functions, we turned to microscopic analysis of BDLP on membranes templates. We find that the GTP-bound BDLP forms a rigid scaffold that is capable of bending and imposing a defined curvature. In contrast, the GDP-bound state is loosely organized and unable to remodel membrane shape. Our results indicate that BDLP intrinsically functions as a protein scaffold whose organization is regulated by nucleotide binding.

## Introduction

Remodeling of membrane shape is fundamental to several cellular processes. The dynamin superfamily of proteins (DSPs) feature prominently in membrane remodeling processes (Ford and Chappie, 2019; Jimah and Hinshaw, 2019). DSPs are peripheral or integral membrane proteins with the former category usually associated with membrane tubulation or fission and the latter with membrane fusion. Soluble dynamins assemble into helical scaffolds. Self-assembly results in stimulated GTP hydrolysis, which drives constriction of the scaffold leading to membrane fission (Schmid and Frolov, 2011; Khurana and Pucadyil, 2023; Sarkar and Pucadyil, 2025). These proteins are characterized by low affinity for GTP, high basal GTPase activity, and the ability to self-assemble into ordered structures that facilitate membrane remodeling (Praefcke and McMahon, 2004; Daumke and Praefcke, 2018; Ford and Chappie, 2019; Jimah and Hinshaw, 2019).

Discoveries in the early 2000s revealed the presence of dynamin-like proteins in bacteria thus expanding the evolutionary significance of this protein family (Bramkamp, 2012). Bacterial dynamin-like proteins (DLPs) share key structural elements with their eukaryotic counterparts, including a conserved GTP-binding domain (G domain), the bundle signaling element (BSE) and the stalk domain. BDLP from the cyanobacterium *Nostoc punctiforme* was the first bacterial DLP to be reported and it appears to be involved in membrane dynamics in cyanobacteria (Low and Löwe, 2006; Jilly *et al*., 2018). Studies on DLPs in other bacteria suggest functions in a range of cellular processes associated with membrane dynamics. Dynamin-like protein A (DynA) from *Bacillus subtilis* has been shown to mediate membrane tethering and fusion in vitro (Bürmann *et al*., 2011; Guo and Bramkamp, 2019). DynA localizes to the membrane in response to stresses like pore formation thereby indicating a role in surveillance (Sawant *et al*., 2016; Guo *et al*., 2022; Sattler and Graumann, 2022). DynA and DynB from *Streptomyces venezuelae* function in a cell division complex required for the formation of the septum during cell division (Schlimpert *et al*., 2017). The *Escherichia coli* DLP LeoA is involved in secretion of toxin-containing vesicles (Brown and Hardwidge, 2007; Michie *et al*., 2014). The *Campylobacter jejuni* DLP form a heterotypic pair and facilitates membrane tethering (Liu *et al*., 2018). In *Mycobacterium smegmatis*, the antibiotic isoniazid induces expression of the *iniBAC* operon, where Isoniazid-induced protein A (IniA) and IniC are DLPs. IniA localizes to membrane ruffles formed upon treatment of cells with isoniazid (INH), an antibiotic that disrupts mycolic acid synthesis and causes membrane stress (Wang *et al*., 2019). More recently, IniA and IniC have been reported to be required for vesicle secretion from mycobacterium (Gupta *et al*., 2023).

The structural organization of Nostoc BDLP is predicted to be like that of the mitochondrial mitofusin (MFN) in the G domain, BSE and stalk (Qi *et al*., 2016; Yan *et al*., 2018). BDLP also shares similarities with the mitochondrial Optic atrophy protein 1 (Opa1) because of membrane binding paddle domain (PD) and the ability to form protein scaffolds (Nyenhuis *et al*., 2023; Von Der Malsburg *et al*., 2023). Both MFN and Opa1 function in membrane fusion and BDLP may function similarly. To address the intrinsic mechanistic attributes of BDLP in membrane remodeling, we evaluated BDLP functions on membrane templates.

## Results

### Membrane binding and GTPase activity of BDLP

Previous studies using vesicle sedimentation assays have reported that the *Nostoc* DLP (BDLP) binds membranes formed with an *E. coli* lipid extract, which contains phospholipids like cardiolipin (CL), phosphatidylglycerol (PG) and phosphatidylethanolamine (PE) (Low and Löwe, 2006; Low *et al*., 2009). Cyanobacteria only contain two anionic phospholipids, phosphatidylglycerol (PG) and the sulfolipid sulfoquinovosyl diacylglycerol (SQDG), along with two uncharged galactolipids monogalactosyl diacylglycerol (MGDG) and digalactosyl diacylglycerol (DGDG) (Omata and Murata, 1983; Mikhaylenko *et al*., 1998). The related *Synechocystis* DLP has been reported to bind SQDG and PG, with PG showing better binding (Gewehr *et al*., 2023). We therefore assayed BDLP functions on PG-containing membranes. Membrane binding was analyzed using Proximity-based Labeling of Membrane Associated Proteins (PLiMAP) (Jose *et al*., 2020). Purified BDLP was incubated with vesicles of increasing PG concentrations. Vesicles contained a small (1 mol%) concentration of a bifunctional lipid, which contains a photoactivable crosslinker moiety at the head and a fluorescent moiety at the tail. Samples were exposed to UV, which cross links the fluorescent lipid to membrane-bound BDLP. The amount of membrane bound BDLP was then quantitated by resolving the sample on an SDS-PAGE and analyzing the in-gel fluorescence associated with BDLP (Fig.1A). These assays revealed robust binding of BDLP above a critical ∼60 mol% PG concentration (Fig. 1A,B). Beyond this critical PG concentration, membrane binding showed a sharp increase with an increase in PG concentration. We therefore chose an intermediate concentration of 80 mol% PG for all subsequent assays.

**Fig. 1.**
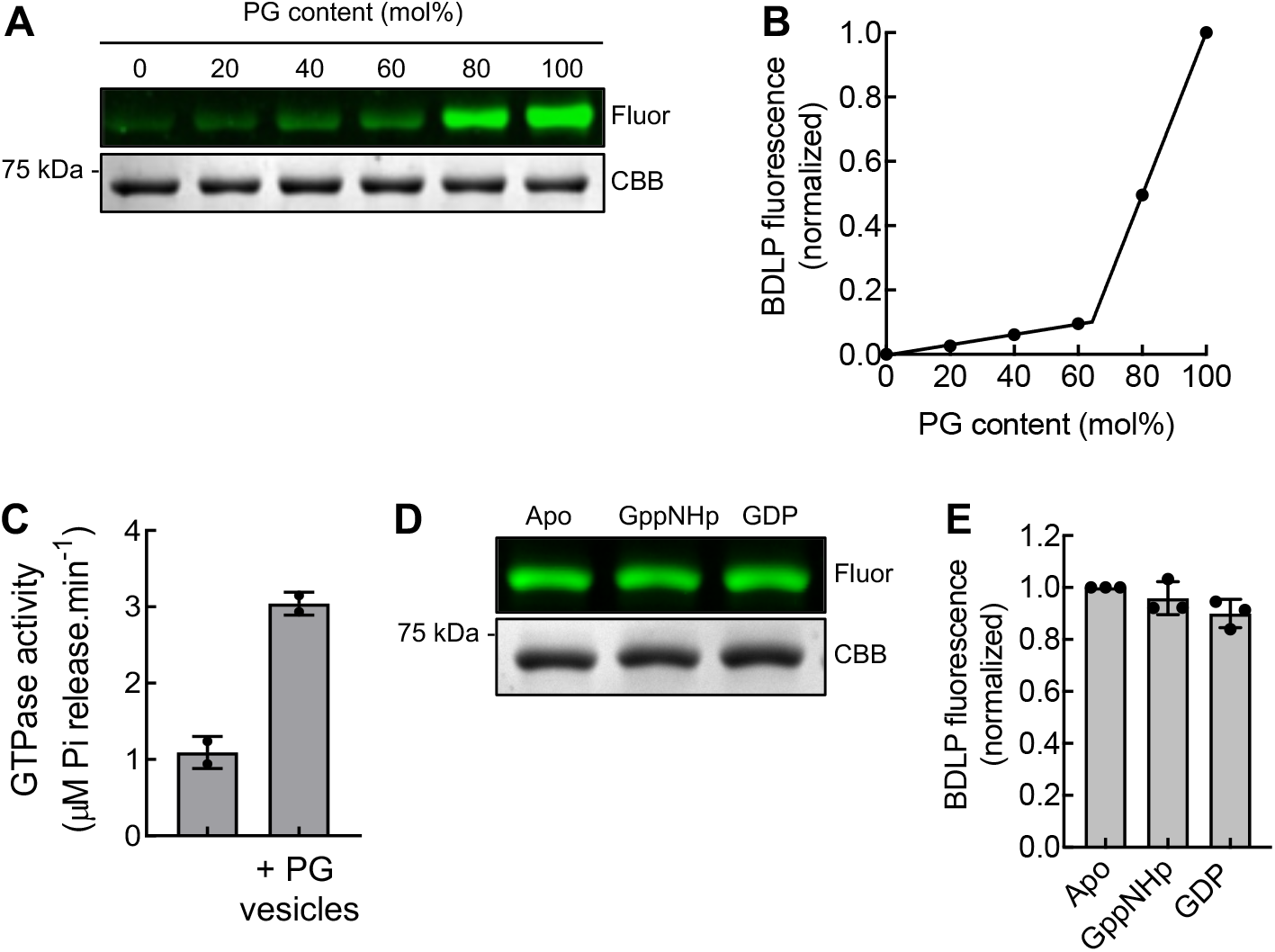
Membrane binding and GTPase activity of BDLP. (A) Results from a representative PLiMAP experiment showing in-gel fluorescence (Fluor) and Coomassie brilliant blue (CBB) staining of BDLP incubated with vesicles containing the indicated concentrations of PG. Experiments contained 1 μM BDLP with 100 μM total lipid. (B) Quantitation of membrane-bound fluorescent BDLP. Data is normalized to fluorescence seen with vesicles containing the highest PG content and represents the mean ± SD of two experiments fitted to a segmental linear regression. (C) GTP hydrolysis activity of 1 μM BDLP alone or in the presence of 80 mol% PG-containing vesicles (100 μM total lipid) (+ PG vesicles). Data represents the mean ± SD of two experiments. (D) Results from a representative PLiMAP experiment showing in-gel fluorescence (Fluor) and Coomassie brilliant blue (CBB) staining of 1 μM BDLP incubated with PG vesicles (100 μM total lipid) in the absence of any nucleotide (Apo) or with GppNHp or GDP. (E) Quantitation of membrane-bound fluorescent BDLP under conditions described in (D). Data is normalized to fluorescence seen in the Apo condition and represents the mean ± SD of 3 experiments.

DLPs display a tendency to self-assemble into helical scaffolds, either at high concentrations in solution or at low concentrations on membrane templates (Ford and Chappie, 2019; Jimah and Hinshaw, 2019). Intermolecular interactions in these assemblies reorient catalytic residues in the G domains for optimal GTP hydrolysis, thereby stimulating their basal rate of GTP hydrolysis of ∼ 1.min^-1^ by 10-100-fold. BDLP displayed a basal GTP hydrolysis rate of ∼ 1.min^-1^, similar to that reported earlier (Low and Löwe, 2006), but showed a modest 3-fold increase with 80% PG-containing vesicles (PG vesicles, Fig. 1C). To assess if membrane binding is nucleotide-dependent, we compared binding in the absence (Apo) or presence of the non-hydrolyzable GTP analog GppNHp and GDP (Fig. 1D). These assays revealed similar levels of membrane-bound BDLP (Fig. 1D,E), indicating that membrane binding of BDLP is nucleotide-independent.

### BDLP-mediated membrane remodeling

BDLPs from *Nostoc* and *Synechocystis* have previously been shown to bind and distort vesicles into narrow tubules (Low *et al*., 2009; Gewehr *et al*., 2023), and the resultant membrane destabilization has been proposed to induce membrane fusion (Low *et al*., 2009; Jimah and Hinshaw, 2019; Gewehr *et al*., 2023; Junglas *et al*., 2024). While membrane fusion has been experimentally demonstrated with the *Synechocystis* DLP (Gewehr *et al*., 2023; Junglas *et al*., 2024), to the best of our knowledge, a similar analysis has not been carried out with the *Nostoc* DLP. We tested if BDLP can fuse PG vesicles using a bulk FRET-based fluorescent lipid dequenching assay (Struck *et al*., 1981). These assays were carried out under conditions like those used for membrane binding and GTPase assays, with BDLP (1 μM) and vesicles (100 μM total lipid). 10 μM PG vesicles containing both the donor fluorescent lipid NBD-PE and the acceptor fluorescent lipid Rh-PE (labeled vesicles) were mixed with 90 μM PG vesicles containing no fluorescent lipid (unlabeled vesicles). Control experiments showed that the donor is efficiently quenched in presence of the acceptor, with a FRET efficiency of ∼80% (Fig. 2). Fusion of the labeled vesicles with excess unlabeled vesicles should dilute the fluorescent lipids in the membrane, thereby lowering this FRET efficiency. Experiments in the presence of BDLP with various nucleotides however showed no change in the FRET efficiency (Fig. 2), indicating that BDLP does not fuse PG vesicles.

**Fig. 2.**
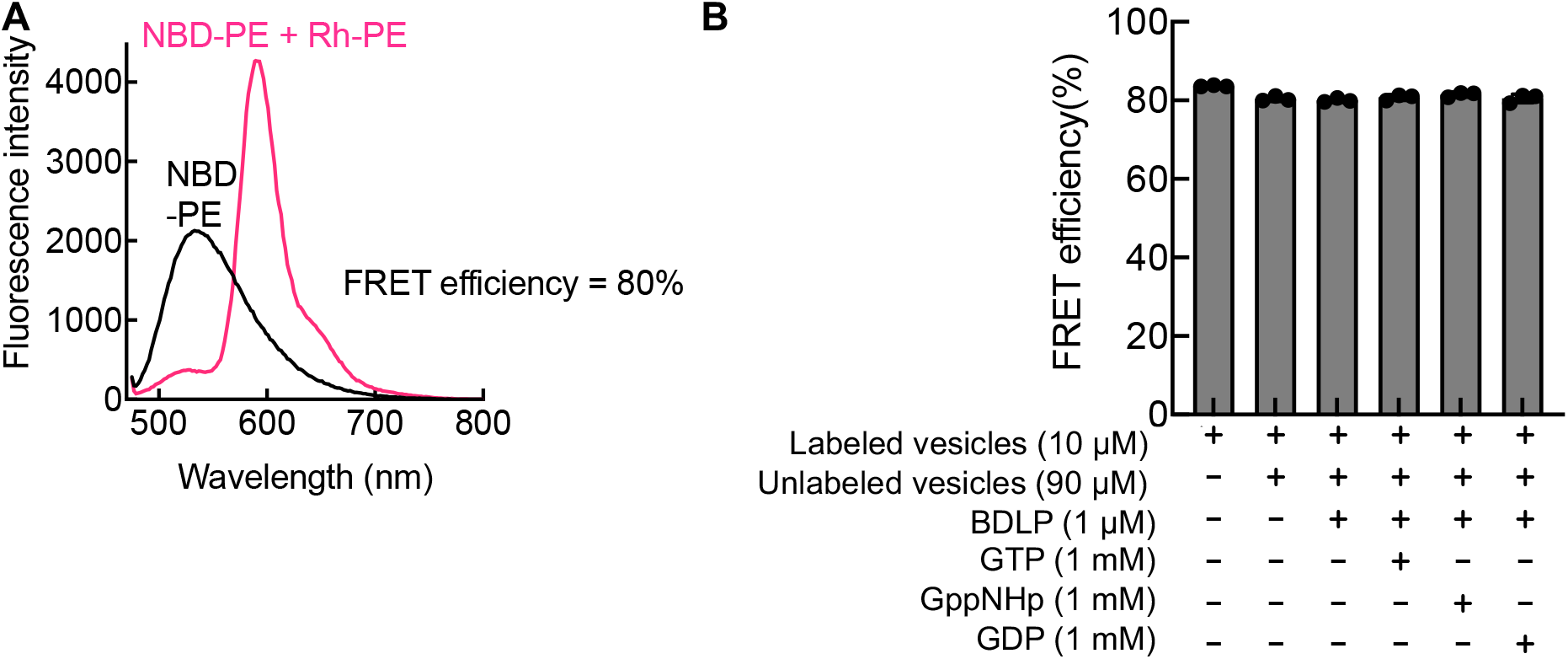
Testing BDLP for membrane fusion. (A) Fluorescence spectra of PG vesicles containing 1 mol% of the donor fluorescent lipid NBD-PE lipid alone (black) or in presence of 1 mol% of the acceptor fluorescent lipid Rh-PE (magenta). The extent of quenching of NBD-PE fluorescence monitored at 535 nm in the presence of Rh-PE is used to calculate the FRET efficiency, which is estimated at 80%. (B) FRET efficiency of PG vesicles estimated in the presence of BDLP and different nucleotides. Data represent the mean ± SD of 3 experiments.

We then turned to a spatially resolved microscopic assay using Supported Membrane Templates (SMrTs) to analyze BDLP function (Dar *et al*., 2017; Bhattacharyya and Pucadyil, 2024; Swaminathan and Pucadyil, 2024). SMrTs display planar lipid bilayers and curved membrane tubes resting on a passive PEG cushion (Fig. 3A). Templates were prepared with a lipid mix containing 80 mol% PG and 1 mol% of the fluorescent lipid Texas Red-DHPE in a DOPC background and displayed characteristic morphologies of planar bilayers and membrane tubes (Fig. 3B). Remarkably, incubating these templates with 0.5 μM of BDLP mixed with 1 mM GTP for 30 mins showed extensive buds (Fig. 3B, red arrowheads) and tubules (Fig. 3B, black arrowheads) on the planar bilayer (Fig. 3A). On the other hand, preformed membrane tubes in SMrT templates appeared striated, with alternating regions of bright and dim fluorescence (Fig. 3B, yellow arrowheads). Reactions with BDLP mixed with GppNHp produced similar effects as those seen with GTP, both on the planar bilayer and on membrane tubes (Fig. 3A). But reactions with BDLP mixed with GDP showed no such effect, with the planar bilayer and membrane tubes appearing just as they did before protein addition (Fig. 3A). Effects on the planar bilayer and membrane tubes appeared largely similar between reactions with GTP or GppNHp, which could arise from the low GTPase activity of membrane-bound BDLP (Fig. 1C, 2B). To understand the mechanistic basis of these effects, we turned to a BDLP construct with a C-terminal mEGFP tag (BDLP-GFP) and carried out experiments in the presence of different nucleotides. Compared to the untagged protein, BDLP-GFP showed a marginally higher basal rate of GTP hydrolysis of ∼2 min^-1^, but a similar rate of ∼3 min^-1^ with PG vesicles (data not shown). BDLP-GFP (1 μM) was mixed with GppNHp and flowed onto the templates. Reactions were incubated for 10 mins and the templates were imaged after washing off excess protein. BDLP localized to both the planar regions and regions that appear to have budded out of the bilayer (Fig. 3C, white arrowheads on the planar bilayer). On membrane tubes, BDLP-GFP was found organized as discrete foci, which coincided with regions of dimmer membrane fluorescence (Fig. 3C, white arrowheads on membrane tubes, Fig. 3D). Since membrane tubes are diffraction-limited objects, regions of dimmer fluorescence reflect thinner or constricted regions on the tube. Dimmer fluorescence could also arise from changes in the fluorescent properties of the lipid probe Texas Red-DHPE or its dilution in the membrane because of an expansion in membrane area upon protein insertion. Indeed, the recently reported structure of the membrane-bound mitochondrial DLP Opa1, which also contains a PD like BDLP, indicates significant displacement of membrane lipids within the plane of the membrane (Nyenhuis *et al*., 2023; Von Der Malsburg *et al*., 2023). To verify this, we used a strategy that was previously applied to analyze the nature of membrane remodeling by other DLPs (Dar and Pucadyil, 2017; Deo *et al*., 2018). We prepared membrane tubes with a 6xHis-tagged GFP tethered to the inner leaflet of the membrane (Fig. 3E). GFP fluorescence is insensitive to its environment and a change in its fluorescence intensity should signify a change in the actual size of the tube. Like was seen earlier (Fig. 3C), addition of BDLP with GppNHp to such tubes displayed regions of low membrane fluorescence, and these regions coincided with low GFP fluorescence (Fig. 3E,F), thus indicating that BDLP foci indeed thin down or constrict the underlying tube. In contrast, reactions with GDP showed BDLP-GFP to be largely uniformly distributed on planar bilayers and membrane tubes with no apparent budding or tubulation of the bilayer or thinning of the tube (Fig. 3E,F).

**Fig. 3.**
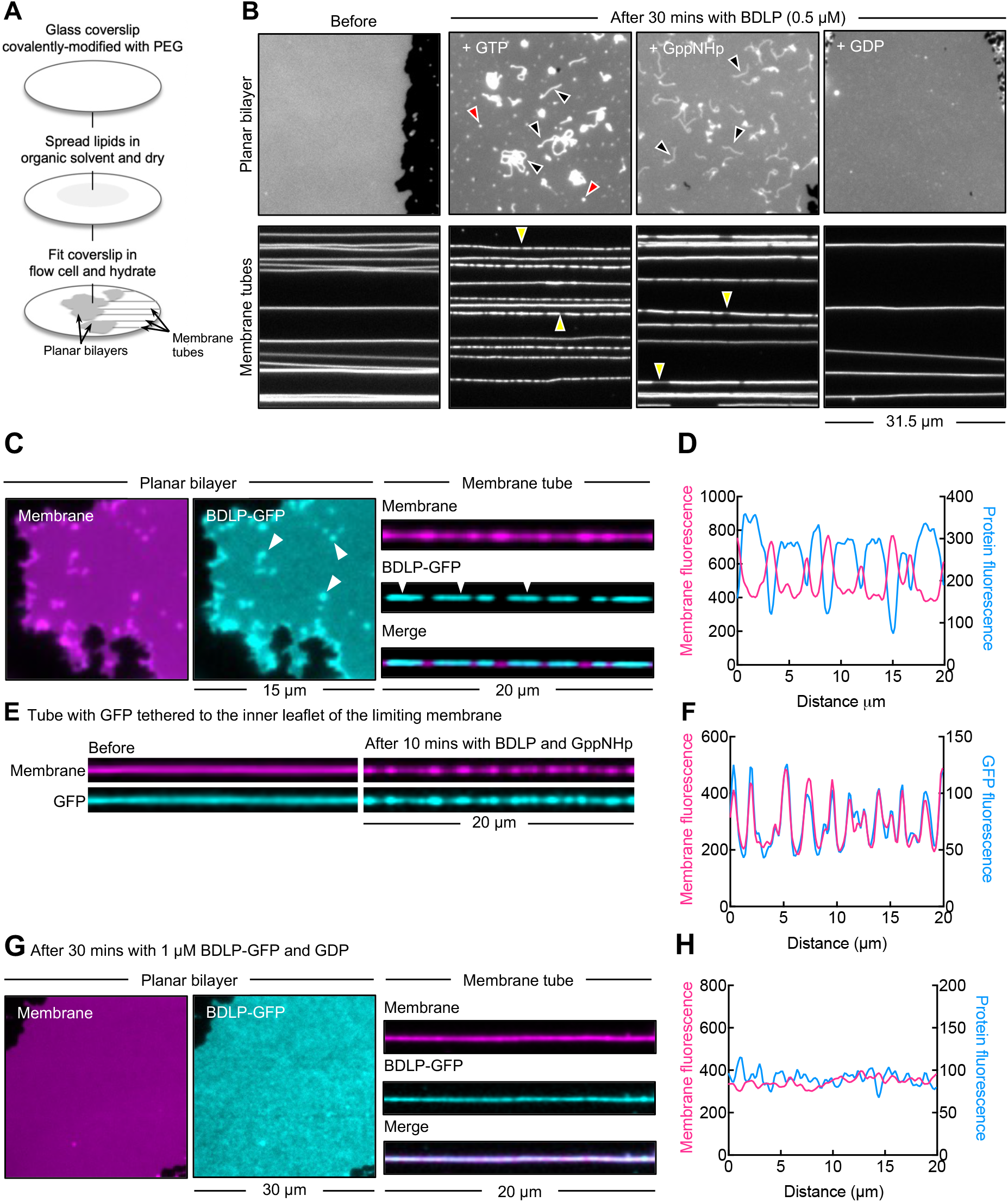
BDLP-mediated membrane remodeling. (A) Schematic showing the preparation of Supported Membrane templates (SMrTs) containing planar bilayers and membrane tubes. (B) Representative images of the planar bilayer and membrane tubes formed of 80% PG before and after 30 mins of incubation with 0.5 μM BDLP with GTP, GppNHp or GDP. Red arrowheads mark membrane buds, black arrowheads mark membrane tubules and yellow arrowheads mark regions of dim tube fluorescence. (C) Representative images of the planar bilayer and membrane tube after 10 mins of incubation with 1 μM BDLP-GFP with GppNHp. White arrowheads mark membrane buds. (D) Profile of membrane and BDLP-GFP fluorescence along the length of the membrane tube shown in (C). (E) Representative fluorescence images of a membrane tube with 6xHis-GFP tethered to the inner leaflet before and after the addition of BDLP with GppNHp. (F) Profile of membrane and GFP fluorescence along the length of the tube before and after BDLP addition. (G) Representative fluorescence images of the planar bilayer and membrane tube after 30 mins of incubation with 1 μM BDLP-GFP with GDP. (H) Profile of membrane and BDLP-GFP fluorescence along the length of the membrane tube shown in (G).

Results from PliMAP assays and reconstitution on SMrT templates indicate that BDLP remains bound to the membrane in both the GppNHp- and GDP-bound states. The GppNHp-bound state but not the GDP-bound BDLP appears to be capable of budding or tubulating the planar bilayer and thinning down membrane tubes. These results could reflect a mechanism whereby GppNHp-binding promotes self-assembly of membrane-bound BDLP into protein scaffolds, which the GDP-bound protein is incapable of. To understand this better, we analyzed tube thinning and membrane organization of different nucleotide-bound states of BDLP.

### Nucleotide-specific organization and dynamics of BDLP

Protein scaffolds usually display an ordered arrangement of protein subunits stabilized by extensive intermolecular interactions with a characteristic and well-defined geometry. To understand if the BDLP assemblies on the membrane are rigid scaffolds that impose a defined curvature, we correlated the size of membrane tubes before and after BDLP addition for tubes of a range of sizes. We acquired images of the same field of membrane tubes before and after BDLP addition, estimated the tube radius using a calibration method and plotted the starting tube radius against that attained after BDLP addition under BDLP foci. If the protein did not constrict the underlying tube, such a plot would show a straight line with a slope of one (Fig. 4A, black dotted line). For GDP-bound BDLP, the slope of such a plot was closer to one (Fig. 4A, blue). However, for GTP-bound (Fig. 4A, black) or GppNHp-bound (Fig. 4A, magenta) BDLP, such a plot revealed a shallow slope with a tube radius that hovered around ∼10 nm, irrespective of the starting tube radius. The GDP-bound BDLP therefore does not alter the size of the underlying tube while GTP-bound BDLP imposes a defined curvature.

**Fig. 4.**
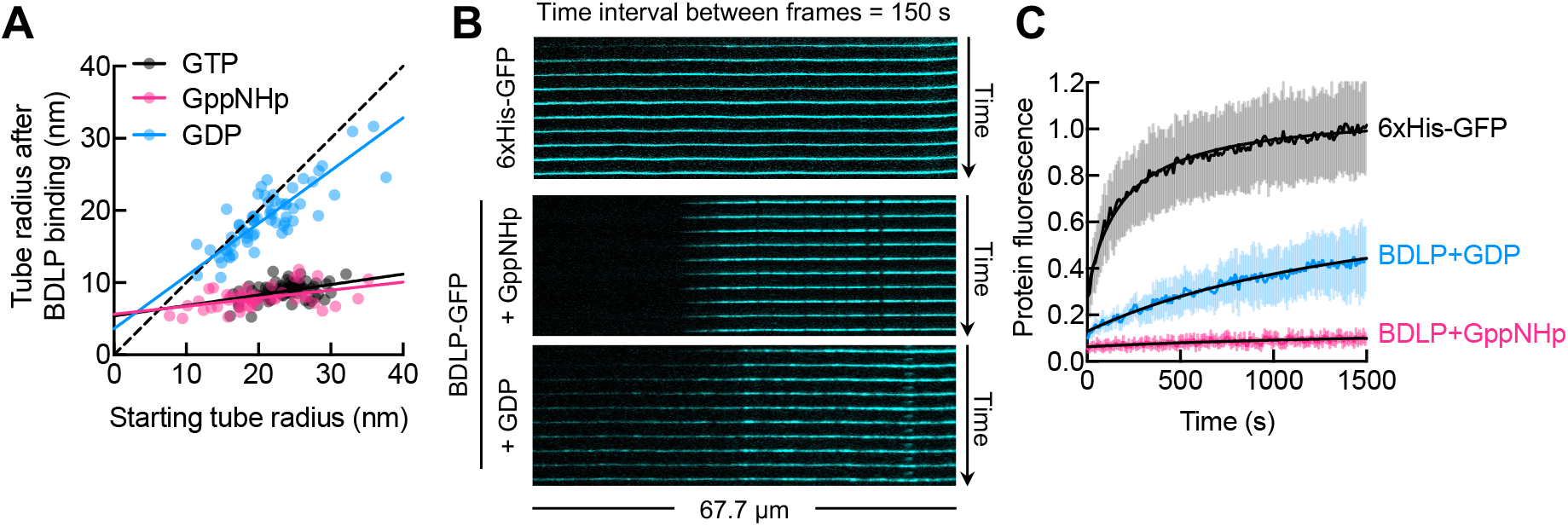
Nucleotide-specific organization and dynamics of BDLP. (A) Plot showing the radius of membrane tubes before and after BDLP addition in presence of GTP (black), GppNHp (magenta) and GDP (blue) across tubes of a range of starting sizes. The black dotted line represents a scenario where protein binding causes no change in tube size. (B) Representative images showing fluorescence recovery after photobleaching of membrane-bound 6xHis-tagged GFP (6xHis-GFP) and BDLP-GFP in the presence of GppNHP or GDP on a membrane tube. (C) Plot showing recovery of fluorescence after photobleaching of the 6xHis-GFP (black) or BDLP-GFP with GppNHp (magenta) or with GDP (blue). Data is normalized to the fluorescence of each construct in a region distant from the site of bleach on the same tube. Data represents the mean ± SD of fluorescence from 13 tubes for 6xHis-GFP, 13 tubes for GppNHp and 12 tubes for GDP, fitted to an empirical fluorescence recovery after photobleaching model (solid black lines).

Scaffolds formed by a peripheral membrane protein like BDLP would be stabilized through a network of intermolecular interactions, which should render them immobile on the membrane. To test this, we monitored the lateral diffusion of BDLP-GFP on membrane tubes by FRAP (Fig. 4B,C). As a mimic of a membrane-bound protein and to establish the timescale for monitoring recovery after photobleaching, we used a 6xHis-GFP and recruited it to the membrane via NTA lipid. The 6xHis-GFP was added to NTA lipid-containing membrane tubes and excess unbound protein was washed off before photobleaching. 6xHis GFP showed full recovery of fluorescence within ∼25 mins (Fig. 4B,C). We then added BDLP-GFP mixed with GppNHp or GDP to membrane tubes, washed off the excess unbound protein and monitored its recovery after photobleaching. Consistent with what is expected of a rigid scaffold, these experiments revealed no recovery for GppNHp-bound BDLP-GFP (Fig. 4C, magenta trace). But the GDP-bound BDLP-GFP showed partial recovery within the same time scale (Fig. 4C, blue trace). The partial fluorescence recovery of GDP-bound BDLP-GFP compared to the NTA lipid-bound 6xHis-GFP could signify differences in how these proteins are bound to the membrane. BDLP binds membrane lipids via the PD, which because of the greater membrane insertion would diffuse more slowly than the NTA lipid-bound 6xHis-GFP. Together, these results signify that GTP-binding promotes self-assembly of membrane bound BDLP into protein scaffolds that can remodel planar bilayers into membrane tubules.

## Discussion

Our results show that the *Nostoc* DLP binds the anionic PG lipid, which is consistent with those reported for the related *Synechocystis* DLP (Gewehr *et al*., 2023). PLiMAP assays reveal that binding is significantly enhanced above a critical PG concentration. The higher binding could reflect an avidity-based increase in affinity for PG that is caused by the self-assembly of membrane-bound BDLP. BDLP binds the membrane through its PD. Although the structure of PD in membrane-bound BDLP is not resolved, previous studies have indicated that mutations in hydrophobic residues in the PD perturb membrane interactions (Low *et al*., 2009). The PD also contains several positively charged residues, which could mediate electrostatic interactions with anionic lipids. Structures of the PD in the mitochondrial Mgm1 and Opa1 emphasize the contribution of both hydrophobic and electrostatic interactions between PD residues and anionic lipids (Low *et al*., 2009; Faelber *et al*., 2019; Nyenhuis *et al*., 2023; Von Der Malsburg *et al*., 2023). Earlier results using vesicle sedimentation assays showed that the GDP-bound BDLP is less prone to sedimentation, which was interpreted to reflect reduced membrane binding (Low *et al*., 2009). But DLPs display a tendency to self-assemble into higher order structures that sediment even in the absence of membranes, which makes it difficult to distinguish between membrane binding and self-assembly from sedimentation assays (Jose *et al*., 2020). Indeed, such assays with the *Synechocystis* DLP in the absence of vesicles show that the GppNHp-bound state is more prone to sediment than the GDP-bound state (Junglas *et al*., 2024). Our results from a more direct PLiMAP assay show that membrane binding of BDLP is nucleotide independent. However, nucleotides influence how BDLP is organized on the membrane. Reconstitution experiments on SMrTs show that the GppNHp-bound BDLP forms immobile assemblies that impose a defined curvature of ∼10 nm radius to the underlying membrane tube. These size estimates are consistent with the cryo-EM structure of BDLP in the GppNHp-bound state (Low *et al*., 2009). On the other hand, the GDP-bound BDLP is mobile, distributes uniformly on the membrane and is incapable of imposing a defined curvature on membrane tubes. These results are consistent with a model wherein GTP binding facilitates self-assembly while GTP hydrolysis causes disassembly of BDLP. The solution structure of the GDP-bound BDLP reveals a closed and bent conformation while the cryo-EM structure of the GTP-bound BDLP on membranes reveals an organized helical scaffold with BDLP in an open and extended conformation (Low and Löwe, 2006; Low *et al*., 2009). Structures of the GTP-bound BDLP in solution or the GDP-bound BDLP on membranes are currently unavailable. The closed and open conformations could reflect structures of the free and membrane-bound states, respectively. Alternatively, they could reflect nucleotide dependent conformational changes in the membrane-bound state. The GDP-bound closed conformation occludes oligomerization interfaces in the stalk region of BDLP which is not the case with the GTP-bound open conformation. Our results, monitoring the organization and dynamics of membrane-bound BDLP, indicate that the structural transition from the closed to open state likely facilitates BDLP to form a protein scaffold. Structures of Mgm1 and Opa1 reveal significant insertion of the PD in the outer leaflet (Faelber *et al*., 2019; Nyenhuis *et al*., 2023; Von Der Malsburg *et al*., 2023). Such insertion could lead to an asymmetric expansion of the outer leaflet, thereby inducing local membrane curvature, which along with the tendency to form scaffolds could explain how the GTP-bound BDLP is capable of tubulating planar bilayers.

Consistent with previous results reported for both the *Nostoc* and *Synechocystis* BDLP (Low and Löwe, 2006; Gewehr *et al*., 2023), membrane binding of BDLP causes a modest ∼3-fold stimulation of its GTP hydrolysis activity. For the endocytic and mitochondrial soluble dynamins involved in fission, membrane binding elicits a robust stimulation of their basal GTP hydrolysis activity, which is a prerequisite for inducing conformational changes in the membrane bound helical scaffolds formed by these proteins for constriction and fission (Dar *et al*., 2015; Kamerkar *et al*., 2018; Ford and Chappie, 2019; Jimah and Hinshaw, 2019). BDLP also forms helical scaffolds and has been speculated to be involved in membrane fission (Low *et al*., 2009), but our results indicate otherwise. The lack of robust stimulation in GTP hydrolysis activity is a characteristic feature among dynamins that are integral membrane proteins such as atlastin and mitofusin, which form GTP-dependent dimers in trans to tether and fuse membranes (Byrnes *et al*., 2013; Cao *et al*., 2017; Li *et al*., 2019). These proteins are not reported to self-assemble into helical scaffolds. On the other hand, the shorter forms of Mgm1 and Opa1 lacking the transmembrane domain form helical scaffolds and show robust stimulation in GTP hydrolysis (Faelber *et al*., 2019; Zhang *et al*., 2020). These proteins are thought to function in membrane fusion by a mechanism that involves membrane destabilization upon tubulation, much like what has been reported for the *Synechocystis* BDLP (Faelber *et al*., 2019; Gewehr *et al*., 2023; Nyenhuis *et al*., 2023; Von Der Malsburg *et al*., 2023). But our results indicate that BDLP does not cause membrane fusion. Our results instead indicate that BDLP forms stable scaffolds that tubulate the membrane in a nucleotide-specific manner. These scaffolds define a membrane curvature of ∼10 nm radius, which would exert extreme curvature stresses on the membrane. Structural analyses of membrane-bound DLPs, both from *Nostoc* and *Synechocystis* appear lacking in the electron density usually associated with the lamellar organization of lipids in the underlying membrane tubule (Low *et al*., 2009; Junglas *et al*., 2024). On the other hand, a lamellar organization of lipids can be clearly seen in the membrane-bound structures of Mgm1 and Opa1 (Faelber *et al*., 2019; Nyenhuis *et al*., 2023; Von Der Malsburg *et al*., 2023). It is possible that the PD in the bacterial DLPs adopts a conformation that induces significant displacement of the outer leaflet lipids thereby forcing the inner leaflet to adopt a highly distorted structure.

Our present results indicate that BDLP forms protein scaffolds in a nucleotide-dependent manner. Based on the pathway to membrane fission seen with other dynamins (Dar *et al*., 2015; Dar and Pucadyil, 2017; Kamerkar *et al*., 2018; Khurana and Pucadyil, 2023; Pemberton *et al*., 2025; Sarkar and Pucadyil, 2025), the inability for BDLP scaffolds to further constrict the underlying tube to cause fission appears to be limited by its low level of self-assembly stimulated GTPase activity. But these scaffolds could undergo further constriction leading to fission in a more native context. The BDLP we have tested here is one among 4 genes in *Nostoc*, all of which are organized in an operon (Bürmann *et al*., 2011; Jilly *et al*., 2018). It is possible that the 4 BDLPs work together. Such collaboration is seen in *Campylobacter jejuni*, where DLP1 and DLP2 are reported to form tetramers with the central DLP2 dimer flanked by two DLP1 proteins. Tetramerization triggers GTPase activity and further oligomerization facilitates membrane tubulation and tethering (Liu *et al*., 2018). The 2-headed dynamin-like protein DynA in *Bacillus subtilis* serves as an extreme example as it represents a fusion of two genes that produces a protein consisting of two distinct subunits, D1 and D2. D1 is essential for membrane binding and fusion, while D2 facilitates the fusion process (Bürmann *et al*., 2011). Thus, the four BDLP gene products in *Nostoc punctiforme* could potentially interact with each other for function and represents an exciting avenue for future research.

## Acknowledgements

K.S. thanks the Council of Scientific and Industrial Research (CSIR) for a graduate student fellowship. T.J.P. thanks the Howard Hughes Medical Institute for an International Research Scholar’s Grant (Grant No. 55008746), the Anusandhan National Research Foundation for a SUPRA Grant (Grant No. SPR12021100014), and the DBT/Wellcome Trust India Alliance for a Team Science Grant (Grant No. IA/TSG/21/1/600245). We thank Harry Low for the BDLP construct. We thank members of the Pucadyil lab for helpful comments on the manuscript.

## Author Contributions

K.S. and T.J.P. conceived of experiments and developed methods. K.S. performed experiments. K.S. and T.J.P. analyzed data. K.S. wrote the first draft of the manuscript which was worked on by T.J.P. T.J.P. acquired financial support for this study.

## Materials and methods

### Constructs and plasmids

BDLP and BDLP fused with a C-terminal GFP were cloned in pET17b with an N-terminal StrepII and C-terminal 6xHis tags. All the clones were confirmed by sequencing.

### Protein purification

Proteins were expressed in BL21(DE3) cells grown in autoinduction medium at 18°C for 36 h. Bacterial cells were pelleted and stored at -40 °C. The frozen bacterial pellet was thawed in 20 mM HEPES pH 7.4, 500 mM NaCl with a protease inhibitor cocktail tablet (Roche) and lysed by sonication in an ice-water bath. Lysate were spun at 30,000 g for 20 min and the supernatant was incubated with HisPur™ Cobalt Resin (Thermo Fischer Scientific). The resin was washed with 20 mM HEPES pH 7.4, 500 mM NaCl and bound protein was eluted with 20 mM HEPES pH 7.4, 500 mM NaCl, 100 mM EDTA. The elution was loaded onto a StrepTrap HP or XT column (GE Lifesciences), washed with 20 mM HEPES pH 7.4, 150 mM NaCl and eluted with the same buffer containing 2.5 mM desthiobiotin or biotin. Proteins were spun at 100,000 g to remove aggregates before use in assays. Protein concentration was estimated from UV absorbance at 280 nm using the molar extinction coefficient predicted by the Expasy ProtParam tool.

### Vesicles preparation

1,2-dioleoyl-*sn*-glycero-3-phosphocholine (DOPC) and 1,2-dioleolyl-*sn*-glycero-3-phosphatidylglycerol (DOPG) were obtained from Avanti Lipids. The bifunctional photoactivable fluorescent lipid probe, BODIPY-diazirine PE was synthesized according to previous protocols (Jose and Pucadyil, 2020; Jose *et al*., 2020). To prepare vesicles, lipids were aliquoted at desired ratios into a glass tube and dried under high vacuum for 3 hours to form a thin lipid film. Following the drying process, deionized water was added to the dried lipid film to achieve a final lipid concentration of 1 mM. The lipid suspension was then hydrated at 50 ºC for 30 mins, during which it was vortexed vigorously to ensure thorough mixing. The hydrated lipids were extruded through 100 nm pore size polycarbonate filters (Whatman).

### Proximity-based labeling of membrane-associated proteins (PLiMAP)

PLiMAP assays were conducted as previously described (Jose *et al*., 2020). Vesicles (100 μM total lipid) composed of increasing concentrations of DOPG, and 1 mol% of BODIPY-diazirine PE were incubated with BDLP (1 μM) in a final volume of 30 μL of 20 mM HEPES pH 7.4, 150 mM NaCl in the absence or presence of nucleotides (1 mM, Jena Biosciences) and MgCl_2_ (1 mM). Samples were incubated in the dark at room temperature for 30 mins and subsequently exposed to UV light (365 nm; UVP crosslinker CL-1000L) at an intensity of 200 mJ/cm^2^ for 1 min. Following UV exposure, samples were mixed with SDS-PAGE sample buffer, boiled, and resolved by SDS-PAGE. Gels were first imaged for BODIPY fluorescence using a Typhoon Biomolecular Imager (Amersham) and subsequently stained with Coomassie Brilliant Blue (CBB) to detect total protein.

### GTPase assay

BDLP (1 μM) was mixed with GTP (1 mM) in the absence or presence of vesicles (100 μM total lipid) in 20 mM HEPES pH 7.4 with 150mM NaCl and 1 mM MgCl_2_ and incubated at 37 °C. Aliquots were taken at regular intervals and quenched with 0.5 mM EDTA. Inorganic phosphate released was assayed with the malachite green reagent (Leonard *et al*., 2005).

### Membrane fusion assay

Membrane fusion assays were performed by monitoring FRET-based dequenching caused by lipid mixing (Struck *et al*., 1981). Labelled vesicles contained 80% DOPG, 17% DOPC, 1% NBD-PE (donor, Avanti Polar Lipids) and 2% Rhodamine-PE (acceptor, Avanti Polar Lipids). Unlabeled vesicles contained 80% DOPG and 20% DOPC. To analyze membrane fusion, 100 μl of a reaction mix was prepared in 20 mM HEPES, pH 7.4 and 150 mM NaCl and contained labelled vesicles (10 μM), unlabeled vesicles (90 μM), 1 μM BDLP, nucleotides (1 mM) and MgCl_2_ (1 mM). Samples were incubated in the dark at 37 °C for an hour. Fluorescence was measured on a plate reader (Tecan Infinite M200 Pro) with an excitation wavelength of 470 nm.

### Supported Membrane Templates (SMrT), fluorescence imaging and image analysis

Supported membrane templates (SMrT) were prepared as described earlier (Dar *et al*., 2017). Briefly, lipids were aliquoted at desired ratios to a final concentration of 1 mM in chloroform. The lipid mixes also contained the fluorescent lipid pTexas-Red DHPE (Thermo Fisher Scientific) at 1 mol% concentration. 2 μl of the lipid mix was spread with a glass syringe on a PEGylated glass coverslip, dried, and assembled inside an FCS2 flow chamber (Bioptechs). The chamber was filled with 20 mM HEPES pH 7.4, 150 mM NaCl and flowed at high rates to form SMrTs. BDLP (0.5 μM) in 20 mM HEPES pH 7.4, 150 mM NaCl was flowed onto the templates and low flow rates and incubated for 30 mins. Templates were imaged after washing off excess BDLP. Tube sizes were estimated based on a calibration procedure as described earlier (Dar *et al*., 2017). For 6xHis-GFP encapsulation, the lipid mix contain 1 mol% DGS-NTA(Ni^2+^) (Avanti Polar Lipids). 6xHis-GFP (1 μM) was mixed in 20 mM HEPES pH 7.4, 150 mM NaCl and flowed at high rates to form SMrTs. This cause the 6xHis-GFP to get tethered on both the inner and outer leaflets of the membrane tube. Templates are then rinsed with 70 mM EDTA pH 7.4 to dissociate the 6xHis-GFP bound to the outer leaflet. Subsequently, 0.5 μM BDLP was flowed onto SMrTs in 20 mM HEPES pH 7.4, 150 mM NaCl and incubated for 30 mins. Templates were imaged after washing off excess BDLP. For FRAP experiments, preformed nanotubes were first incubated with the desired proteins. Excess protein was washed off and the protein in the tubes was bleached by increasing the incident light intensity. This bleached a large area of the membrane tubes. The incident light intensity was lowered, and the bleached region was imaged at 10 s intervals to monitor recovery of fluorescence. Imaging was carried out through a 100x, 1.4 NA oil-immersion objective on an Olympus IX83 inverted microscope connected to an LED light source (CoolLED) and an Evolve 512 EMCCD camera (Photometrics). Image acquisition was controlled by μManager and analyzed using Fiji.

